# White matter conduction in the human brain is mostly slow, with rare high velocity connections

**DOI:** 10.64898/2026.07.28.741279

**Authors:** Samuel Romero-Santiago, María Guadalupe Yáñez-Ramos, Jordan A. Bilderbeek, Nicholas M. Gregg, Myung-Ho In, Erin Gray, Daehun Kang, Yunhong Shu, Gregory A. Worrell, Cameron C. Mcintyre, Kai J. Miller, Dora Hermes

## Abstract

White matter bundles play a crucial role in cognitive functions by rapidly transmitting information between brain regions. Inter-areal conduction guides the integration of information and conduction velocity is a fundamental parameter in theories and models of brain function. However, distributions of conduction velocity remain difficult to characterize in vivo in humans. In this work, we integrated diffusion magnetic resonance imaging (dMRI) tractography with intracranial electrical stimulation during clinical stereo-electroencephalography (sEEG) monitoring in 17 subjects to measure conduction velocity within four major white matter bundles. Our findings reveal that human brain conduction is characterized by high variability both within and between bundles, reflecting a predominance of slow connections alongside rare high-speed connections. Because conduction velocity in myelinated fibers follows an approximately linear relationship with axon diameter, we derive underlying axon diameter distributions and show that these estimates are comparable to previous post-mortem studies. These findings demonstrate a heavily skewed distribution of human neural conduction velocities and show that structural heterogeneity shapes the timing and integration of information in large-scale networks.

## Introduction

Brain function depends on rapid signal propagation along white matter bundles. While the speed of action potential propagation has been proposed to positively predict cognitive performance [1–3], current models of human brain function do not agree on typical conduction velocities. Some models use an average conduction velocity of 4 m/s while others use 26 m/s [4–10]. Accurately modeling human brain function therefore warrants a detailed investigation of white matter conduction velocity.

Conduction velocities in the human brain are typically estimated from axon diameter, given the approximately linear relationship between axon diameter and conduction velocity [11–16]. Diffusion MRI (dMRI) based microstructural models have estimated axon diameter distributions [1,17,18] and derived conduction velocities [19,20]. However, these estimates depend on biophysical assumptions and potential limitations from small diameter fiber sensitivity and voxel level signal averaging, and therefore require validation. Post-mortem histological evidence across select bundles demonstrates significant variability in axon diameters, with a right-skewed distribution containing many small axons and fewer larger ones [21–25]. Anatomical studies therefore predict that, rather than an average conduction velocity along white matter pathways, conduction velocities should show a right-skewed distribution.

To test this prediction in living human brains, we employ a novel multimodal approach, combining single pulse electrical stimulation during stereotactic EEG (sEEG) monitoring and dMRI based tractography across 17 subjects. Stereo EEG electrodes are placed directly in the white matter allowing direct axonal stimulation, extending previous cortico-cortical approaches with surface electrodes that selectively sample and stimulate gray matter [26,27]. We stimulated four major white matter bundles: the Arcuate Fasciculus (AF), the Uncinate Fasciculus (UF), the Corticospinal Tract (CST) and the Cingulum (Cing). Our results show distinct conduction velocity distributions across these pathways, estimates of underlying axon diameter distributions and a comparison with basic MRI metrics. These distributions reveal that, while all conduction velocity estimates used in current models of human brain function are plausible, they only represent a tiny portion of the full distribution. The full distribution of axon diameters shape conduction velocities and therefore the timing of information integration in large-scale networks.

## Results

To characterize the distribution of conduction velocities in the human brain, we stimulated sEEG electrodes placed in the white matter with a single pulse and measured the latency of evoked responses. In an example subject, Figure 1 shows how an electrode in the mid-segment of the arcuate fasciculus was stimulated and a response was measured at a posterior endpoint of the arcuate. The arcuate fasciculus was segmented from dMRI whole-brain tractography, streamlines connecting these two sites were identified, and the distance along the shortest streamline was 68 mm. The earliest peak in the brain stimulation evoked potential (BSEP) was observed at 5.62 ms. We assumed that the signals travel largely antidromically and therefore synaptic times are negligible [28–32]. The earliest peak is therefore assumed to represent the average of the earliest arrival times of signals at the arcuate endpoint. Therefore, an estimate of the conduction velocity along this streamline is 12.08 mm/ms.

**Figure 1.**
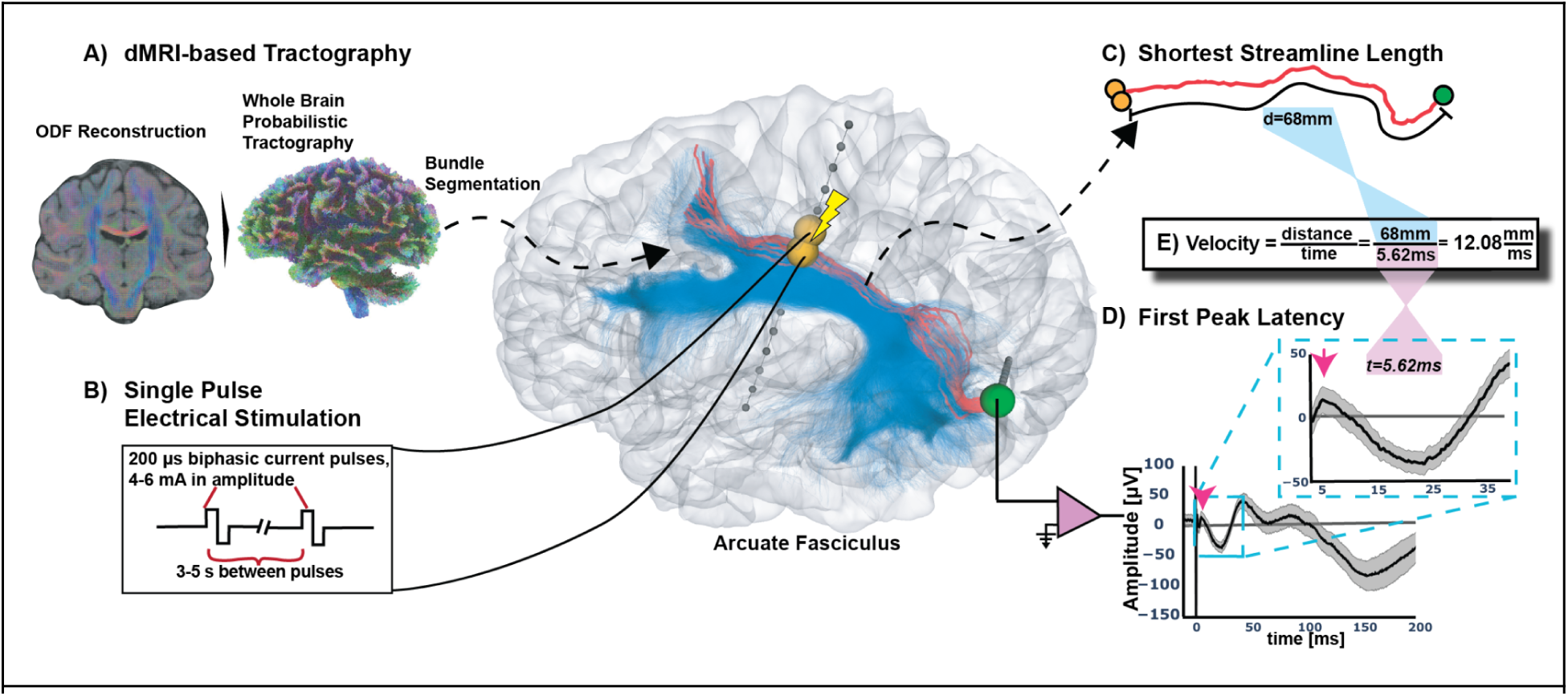
Integration of dMRI and single pulse stimulation to measure Conduction Velocity. **A)** Reconstruction of the Orientation Distribution Function (ODF) and whole-brain probabilistic tractography derived from dMRI in subject 12. Segmentation of the left arcuate fasciculus tractography is shown in blue. **B)** Single pulse biphasic electrical stimulation was delivered through a pair of electrodes (orange spheres) located in the mid-segment of the arcuate fasciculus. **C)** Streamlines connecting the stimulated electrodes with the recorded electrode (green sphere), located at a posterior endpoint of the arcuate fasciculus, were identified (red streamlines). For each streamline, the distances from the stimulated and recorded electrodes were measured, and the conduction distance was defined as the shortest length. **D)** The Brain Stimulation Evoked Potential (BSEP) is shown as a black trace, the latency of the first peak in the response was then identified and considered as the average conduction delay. **E)** Conduction velocity was computed as distance over time.

### Subject-level variability in conduction velocity profiles

At the subject-specific level, analysis was limited by the clinically driven individualized electrode coverage, with most of the subjects having coverage in only a subset of the four bundles. However, for some subjects with denser sampling at endpoints of all 4 tracts, we found high variability in conduction velocities within bundles (Figure 2). Furthermore, visual inspection revealed that the shapes of the velocity distributions were generally comparable across subjects for the same bundle, while remaining distinct between different bundles. The peaks of the velocity distributions suggest a hierarchical organization of conduction velocities. The uncinate fasciculus was consistently identified as the slowest bundle, followed by the cingulum and the arcuate fasciculus with intermediate velocity profiles, while the cortico-spinal tract exhibited the highest conduction velocities across all sampled subjects. Moreover, each bundle shows a high-variance velocity distribution of velocities that is often right skewed.

**Figure 2.**
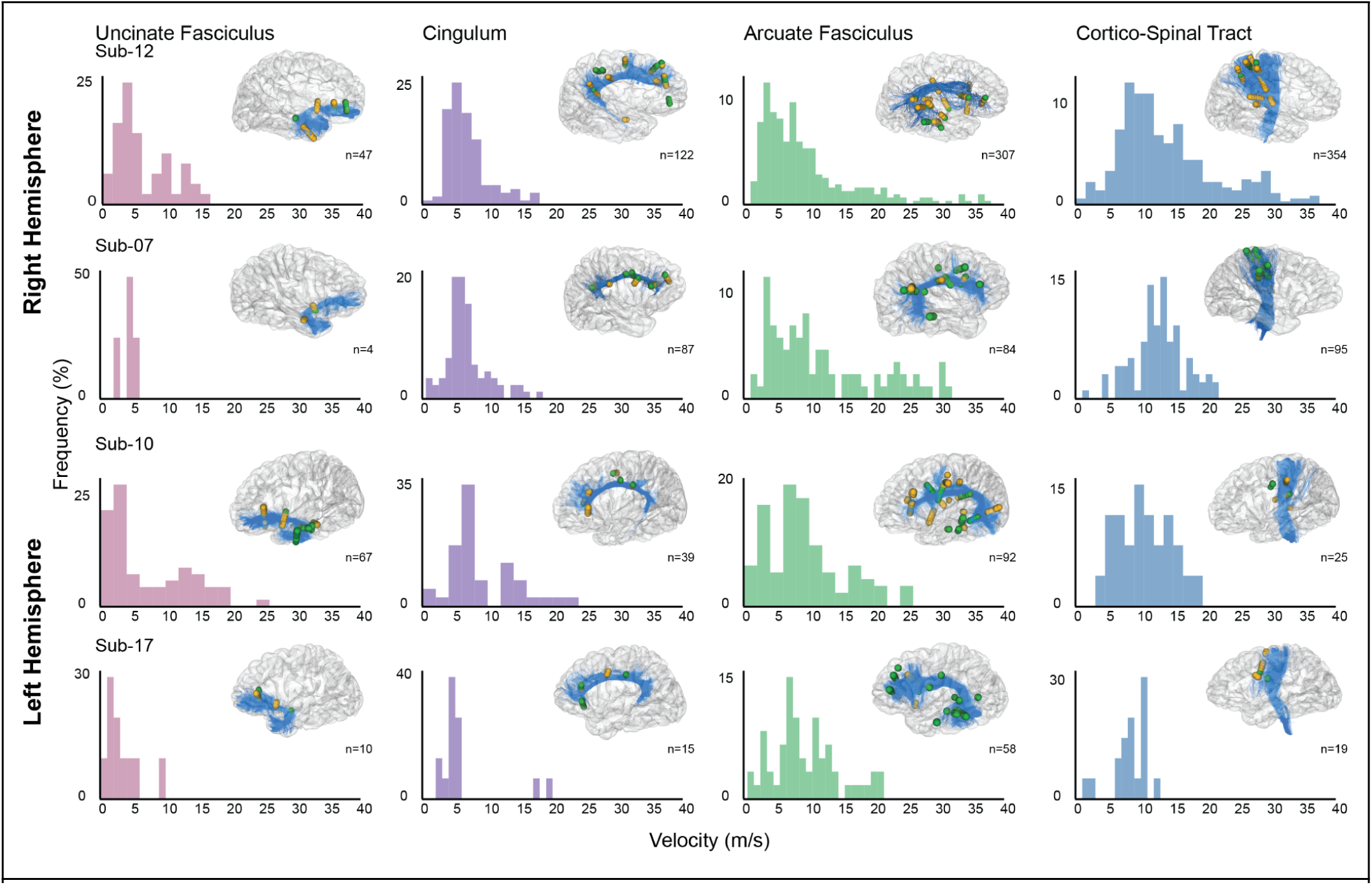
Subject-specific conduction velocity histograms. Each row corresponds to a different subject with electrode coverage in at least one hemisphere across four white matter bundles. Subjects Sub-12 and Sub-07 represent the right hemisphere, while Sub-10 and Sub-17 represent the left. Each column represents a different bundle. Histograms show the distribution of conduction velocities for each bundle. Electrodes with significant responses are shown as spheres within their respective bundle in individual glass-brain renders; orange spheres indicate stimulated electrodes, and green spheres indicate recorded electrodes with significant responses (some orange electrodes were both stimulated and recorded). The number of significant responses (n) within each bundle is reported below each rendering.

### Conduction velocity distributions are long-tailed across subjects and hemispheres

To characterize conduction velocity distribution across participants, the data were combined for each bundle and hemisphere by pooling observations. In Figure 3, distributions for each bundle and hemisphere are shown. All bundles exhibit high variability and a clear long-tail, with high positive skewness (>0.5) observed for most bundles (see Supplementary Table 1 for statistical descriptors). This shows that while most conduction velocities are slow, a few are notably high.

**Figure 3.**
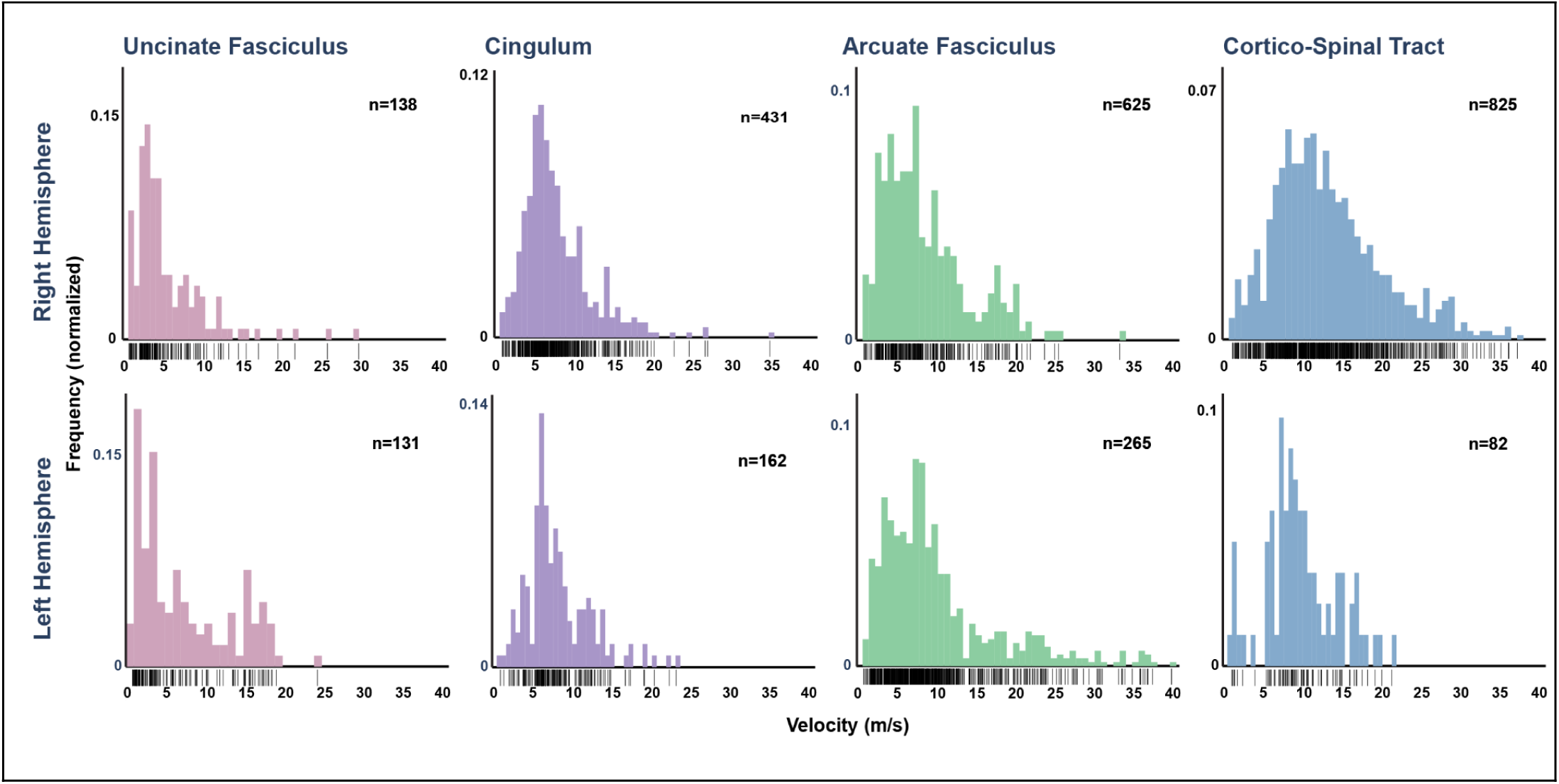
Hemisphere-wise distributions of conduction velocities across white matter bundles and subjects. Pooled velocity estimations are shown as histograms for each bundle (columns), separated by hemisphere (rows). Individual observations are displayed below each histogram as black tick marks. Distributions generally showed positive skewness, with most observations concentrated at lower velocities and a smaller number of higher-velocity observations extending the right tail. No significant left–right hemisphere differences were observed for the uncinate fasciculus, arcuate fasciculus and cingulum (p > 0.05). In contrast, a small but significant hemispheric difference was found for the cortico-spinal tract (p <0.001).

To assess hemispheric lateralization of conduction velocity distributions, group medians were compared using two-sided permutation tests [33] with corrections for multiple comparison. Results show no significant inter-hemispheric differences for the uncinate fasciculus, arcuate fasciculus, or cingulum (p_corrected_<0.05). The right cortico-spinal tract (median 12.2 m/s) showed a small, but significant difference compared to the left (median 9.3 m/s) (p_corrected_<0.001), this result should be interpreted cautiously because of the severe sample size asymmetry between hemispheres (n=825 right vs. n=82 left). Given the uneven sampling and lack of consistent lateralization across bundles, hemispheres were pooled for subsequent analyses.

### White matter bundles exhibit distinct conduction velocity profiles

To assess inter-bundle differences, conduction velocity data were pooled across all subjects and hemispheres for each specific bundle. The resulting histograms are shown in Figure 4, where the cortico-spinal tract exhibited the highest conduction velocities (mean = 13.11 m/s), followed by the arcuate fasciculus (mean = 9.79 m/s) and the cingulum (mean = 8.06 m/s), whereas the uncinate fasciculus (mean = 6.44 m/s) showed the lowest. Mean conduction velocity differed across the bundles (p < 0.001, Welch’s ANOVA). Subsequent pairwise comparisons via the Games–Howell post hoc test revealed significant differences for all bundle pairings (Figure 4, top-right).

**Figure 4.**
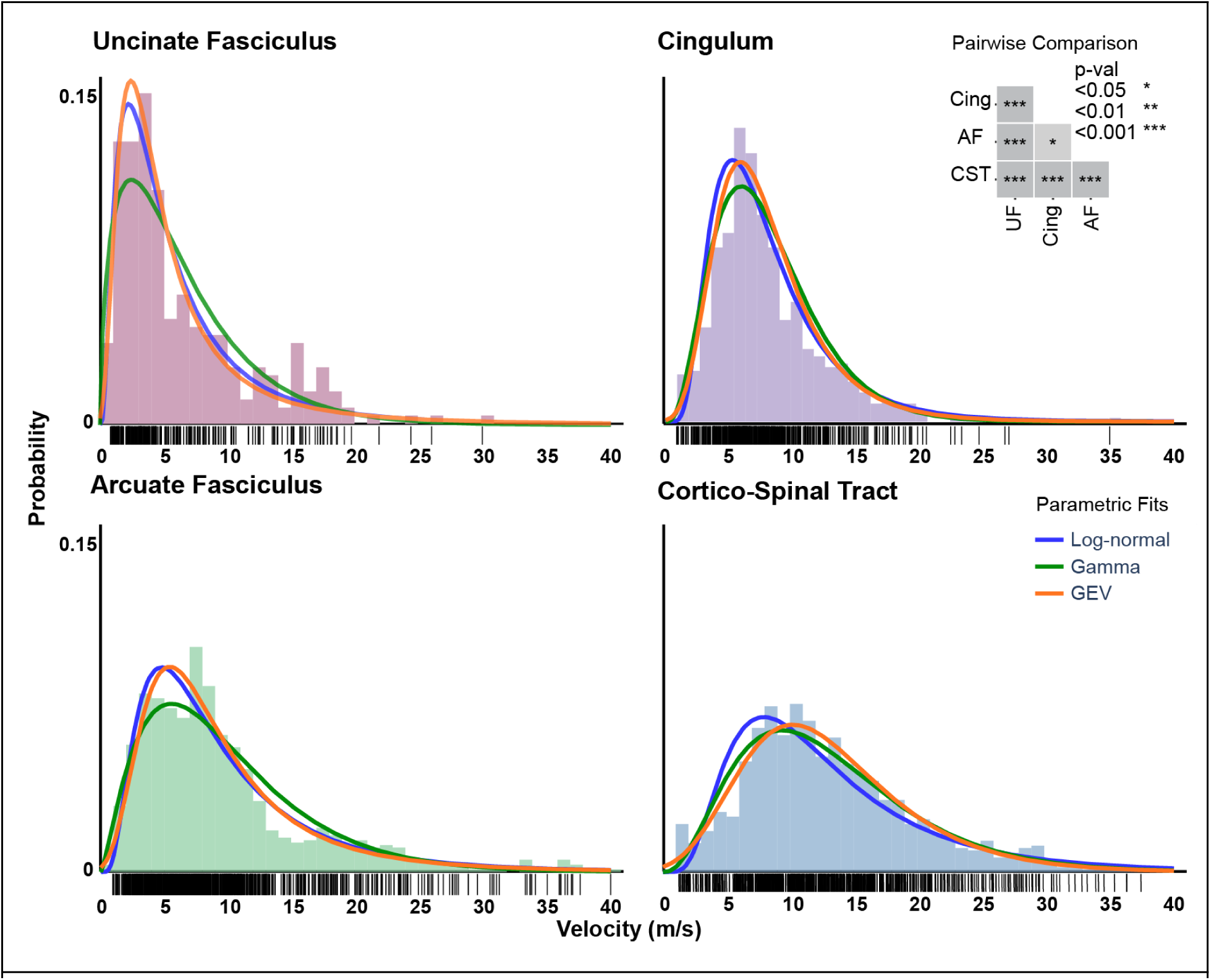
Conduction velocity distribution histograms pooled for four major white matter bundles. Pooled data of all subjects and hemispheres for Uncinate Fasciculus (UF, pink), Arcuate Fasciculus (AF, green), Cingulum (Cing, purple) and Cortico-Spinal Tract (CST, blue). Individual observations are displayed below the histograms as rug plots. Pairwise comparisons via Games-Howell test revealed significant differences between all bundles; AF:CST (p < 0.001), AF:Cing (p = 0.013), AF:UF (p < 0.001), CST:Cing (p < 0.001), CST:UF (p < 0.001), and UF:Cing (p < 0.001). Fitted Log-normal (blue line), Gamma (green line), and Generalized Extreme-Value (orange line) functions are shown for each bundle.

Since many models incorporate conduction velocity or axon diameter as a crucial parameter, it is beneficial to identify a parametric distribution function to properly describe the data. The Gamma and Log-normal distributions are frequently employed for this purpose, as their non-negativity and right-skewed profiles align well with physiological observations [34]. Additionally, the Generalized Extreme Value (GEV) distribution has been proposed as a compelling alternative, demonstrating strong performance in describing axon diameter variations [35,36]. To evaluate the goodness of fit among these three candidates, each model was fitted to the pooled data and assessed using log-likelihood, the Akaike Information Criterion (AIC), and the Bayesian Information Criterion (BIC). These results are summarized in Supplementary Table 2, with the corresponding fits visualized in Figure 4. Analysis of the AIC and BIC indicates that the Log-normal distribution provided the superior fit for the uncinate and arcuate fasciculi, while the GEV distribution performed best for the cingulum and the cortico-spinal tract. For reference, the Log-normal distribution shows that the cortico-spinal tract has the fastest distribution (μ = 2.4237, σ = 0.5918), followed by the arcuate fasciculus (μ = 2.0503, σ = 0.6956) and the cingulum (μ = 1.9527, σ = 0.5377), with the uncinate fasciculus being the slowest (μ = 1.5242, σ = 0.8466).

### Velocity-derived axon diameter estimates provide a qualitative comparison with histology

For each bundle, inner axon diameter distributions were derived based on the reported linear relation with conduction velocity [14]. In myelinated axons, this relationship has the following form:

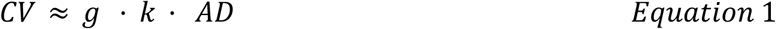

Where CV is the conduction velocity, AD the inner axon diameter, g is a g-ratio of 0.7 [37,38] and k is a constant linear factor that encompasses physiological and structural characteristics, internodal length and biochemical factors [11–16]. Because reported scaling factors vary across studies and tissue conditions, we used Equation 1 with three scaling factors *k* = 4, 6, and 8 based on previous estimates (see methods).

Using this linear relationship, we derived estimates of the inner axon diameter (Figure 5). For comparison, we show previous post-mortem histological data in the uncinate and arcuate fasciculus [22]. A qualitative visual comparison indicates that our results are relatively consistent with these previous post-mortem histological data (frequency polygons adapted from the histograms in [22]). However, our results estimate slightly larger diameters compared to histology, especially in the arcuate fasciculus. This is consistent with the inherent flaws of histological processing, which frequently includes tissue shrinkage resulting from fixation [39–41]. This shrinkage would result in reported diameters that are smaller than their true physiological values, effectively shifting the distribution to the left. Our slightly larger estimates therefore likely provide a close approximation to the underlying anatomical measurements for these two bundles. This comparison suggests that different bundles in the human brain have distinct structural properties that influence the speed of brain dynamics.

**Figure 5.**
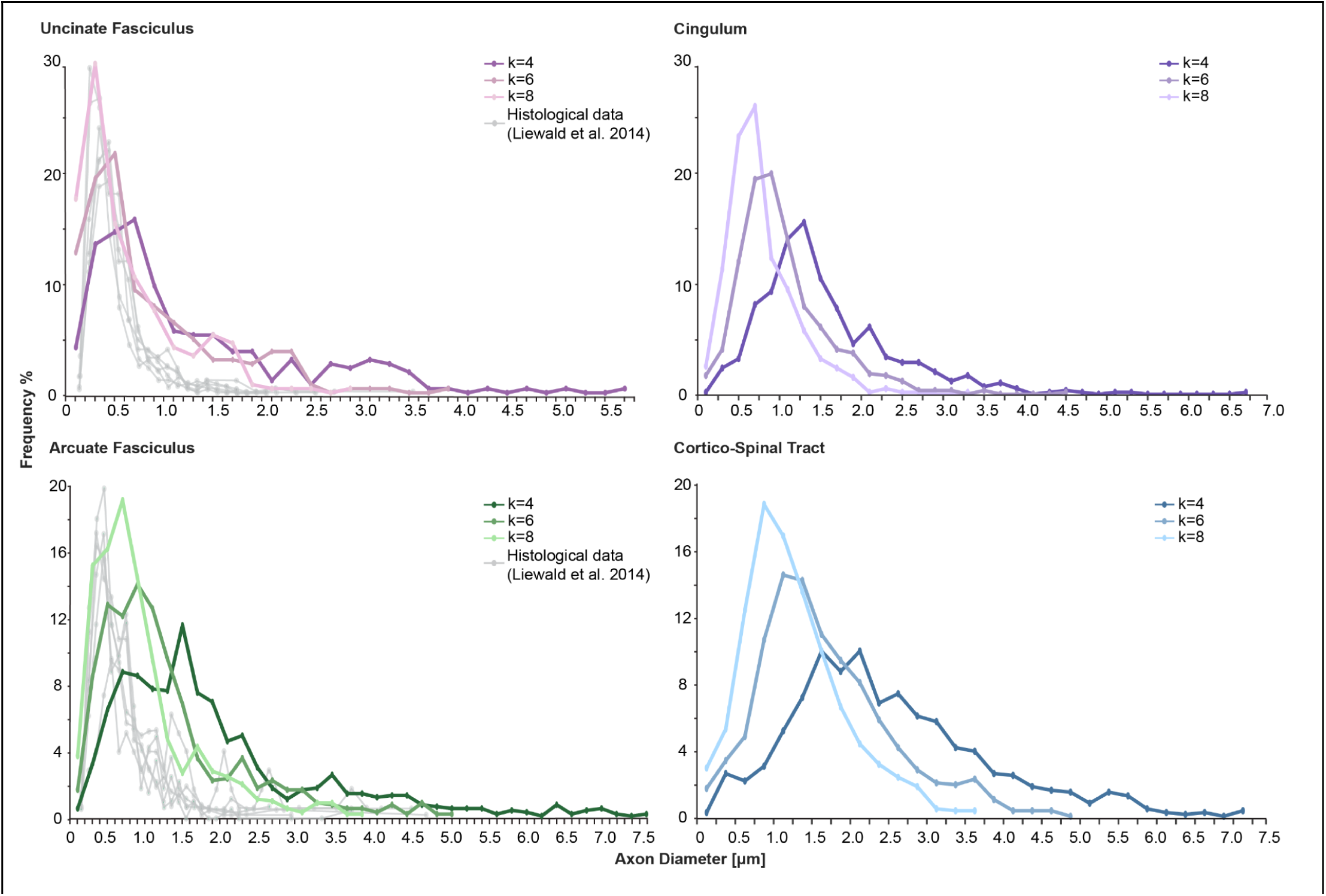
Frequency polygons of estimated inner axon diameters for the selected white matter bundles. Data are shown for the uncinate fasciculus (pink), cingulum (purple), arcuate fasciculus (green) and cortico-spinal tract (blue). After estimating outer diameters using three different linear scaling factors (k = 4, 6, and 8), we applied a constant g-ratio of 0.7 to convert these estimates to inner diameters. For the uncinate and the arcuate fasciculi, histological frequency polygons for inner axon diameters (gray lines) were retrieved manually from [22].

### MRI-derived microstructural metrics are weakly associated with conduction velocity

It is important to understand to what extent the velocity estimates can be explained by MRI-derived microstructural parameters. Diffusion Tensor Imaging (DTI) metrics, specifically Fractional Anisotropy (FA) and Radial Diffusivity (RD), are frequently employed as quantitative descriptors of tissue microstructural properties. FA is commonly interpreted as a measure for the degree of myelination and axonal integrity [42], with higher values correlating with greater myelin content. RD, on the other hand, represents the diffusion perpendicular to the principal diffusion direction. It has also been used as an indirect marker of fiber coherence and myelination, generally showing an inverse relationship with the degree of myelination [42,43]. Another broadly used metric is the T1w/T2w ratio, which was initially proposed as a proxy for myelin content in cortical areas, with higher values representing greater myelination [44–46].

Therefore, we test whether MRI-derived metrics are associated with the conduction velocity measurements. For each connection, we calculate the mean FA, RD and T1w/T2w ratio along the shortest streamline and use this as a predictor of conduction velocity in a Linear Mixed-Effects Model (LMM). The LMM allows us to estimate population-level effects while accounting for between-subject variability. Our results show weak, but statistically significant positive relations between FA and conduction velocity across all white matter bundles (p < 0.05, Wald z-test, Figure 6A, supplementary Table 3). This relation may reflect a degree of covarying fiber coherence and other microstructural properties [42].

**Figure 6.**
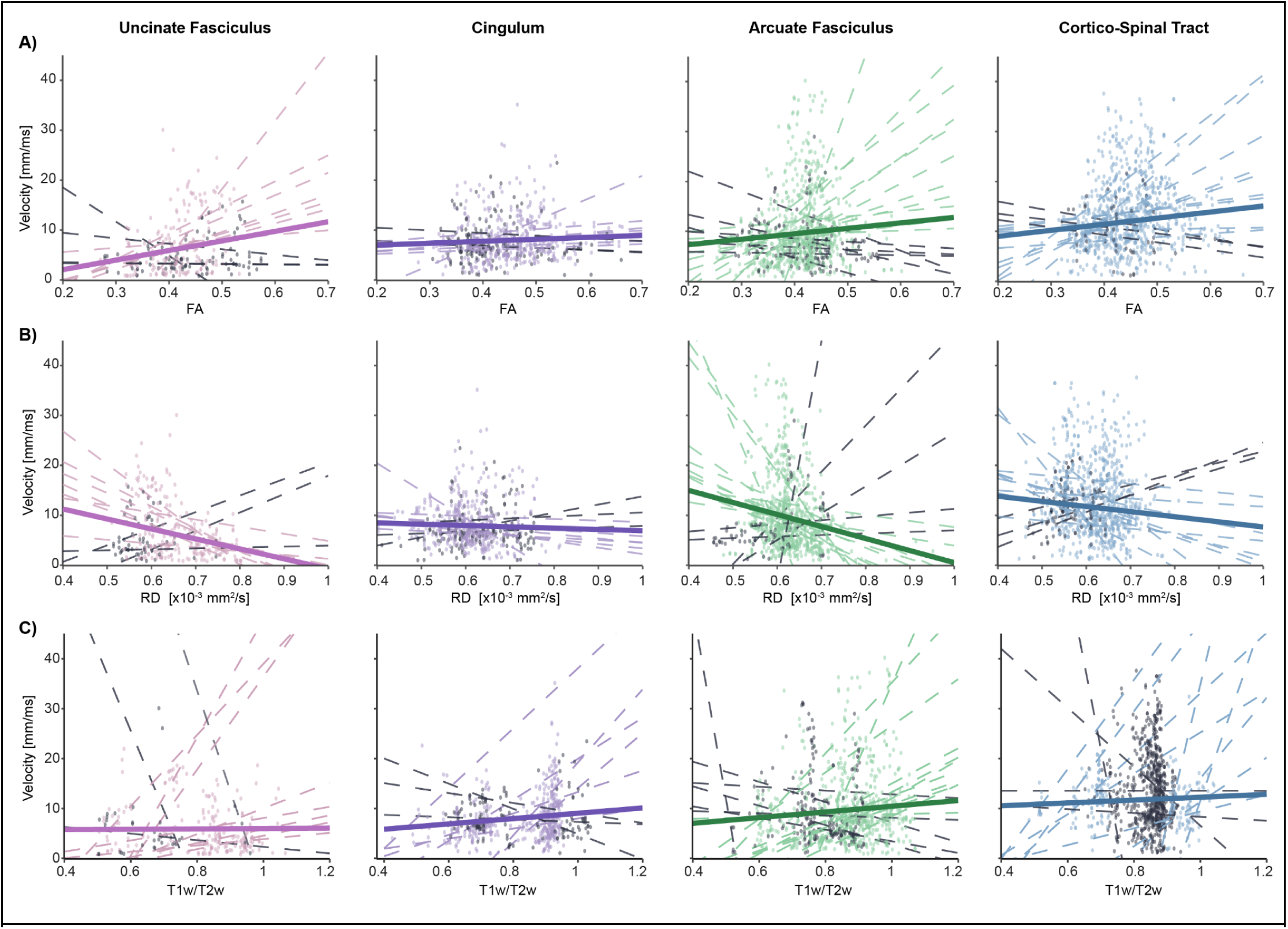
Relationship between MRI metrics and estimated conduction velocities across bundles. **A)** Scatter plots show the relation between fractional anisotropy (FA) and velocity across all measured connections. The thick solid line represents the population-level fit estimated via a Linear Mixed-Effects Model, which shows a positive relationship between FA and velocity. **B)** Scatter plots show the relation between radial diffusivity (RD) and velocity across all measured connections. The population-level fits show a negative relationship for RD and velocity. **C)** Scatter plot shows the relationship between T1w/T2w ratio and conduction velocity across all measured connections, the thick solid line represents the population fit which slope is positive for all bundles. Only for visualization purposes all plots, dashed-lines represent subject-specific Ordinary Least Squares fits, the colored dashed lines indicate the fits for subjects with the same slope sign as the population-level fit whereas black indicates the subjects with slopes of the opposite sign. Individual dots follow the same color code as their corresponding subject-specific fits.

RD demonstrated weak, but significant negative relations with conduction velocity (p < 0.05, Wald z-test) with the exception of the cingulum (p = 0.072) (Figure 6B, supplementary Table 3). Given that reduced RD has been linked to increased myelination [43,47], this finding is consistent with a relationship between conduction velocity and myelination. However, the slope for the cingulum did not reach statistical significance, likely reflecting its complex anatomical organization. Rather than being a unitary pathway, the cingulum comprises short and long sagittal association fibers, including numerous short corticocortical connections linking medial regions of the frontal, parietal, and temporal lobes, as well as fibers radiating toward cortical and subcortical targets along the length of the tract [48]. This heterogeneous fiber organization most likely leads to crossing, bending and fanning fibers which DTI fails to resolve as it only gives a single principal diffusion direction per voxel [49].

Finally, the T1w/T2w ratio showed positive relationship with conduction velocity for all bundles, however, the uncinate fasciculus (p = 0.923, Wald z-test) and the cortico-spinal tract (p = 0.275, Wald z-test) did not reach statistical significance (Figure 6C, Supplementary Table 3). This is not surprising given that this technique was developed to map myelin content in cortical areas [44], while in this work, we mapped white matter structures where the relation has been reported to be limited for deep white matter areas [46,50,51]. While these relations were weak and not consistently observed in each individual subject (Figure 6, dashed-lines), it is promising that the MRI metrics across the population showed the expected associations with conduction velocity.

## Discussion

To approximate axon diameter distributions in vivo from physiological conduction estimates in the human brain, we proposed a novel pipeline that measures conduction velocity by combining sEEG-based electrical stimulation to obtain conduction delay with dMRI-based tractography to determine conduction distance. Axon diameter was then inferred from the estimated conduction velocity. Our findings indicate that across four different fiber bundles tested, conduction velocities showed right-skewed distributions, with many slower connections and only few high speed connections.

### Necessary assumptions to estimate conduction velocity

To estimate conduction velocities in the human brain in vivo, we made some necessary assumptions about the measured latencies of the evoked response. First, we interpret the earliest detectable peak of the evoked response as predominantly reflecting antidromic responses. Direct white matter stimulation triggers bidirectional propagation (both orthodromic and antidromic conduction [52,53]). If there is a direct connection, the first peak is thought to reflect the activation of the fastest-conducting axons within the stimulated fiber population [54]. The first peak of the antidromic response is minimally influenced by synaptic transmission and is expected to provide the closest approximation of axonal conduction delay. Subsequent peaks are interpreted as orthodromic responses, which are typically delayed relative to antidromic activity due to synaptic transmission, on the order of ∼0.5–2 ms [28,31,32,55]. Later components may also reflect the recruitment of polysynaptic responses triggered by the initial input or overlapping delayed signals from the activation of thinner, slower axons. Second, a necessary assumption for this analysis is that there is a direct connection between the stimulated and measured sites. We therefore investigated responses in sites that are connected to the stimulated sites based on dMRI bundle estimates. The anatomical validity of reconstructed bundles strongly depends on image quality, and ambiguities between diffusion orientations and underlying fiber disposition can arise, particularly in regions containing crossing or kissing fibers, where tractography algorithms may produce spurious streamlines and false-positive connections [56]. While these two assumptions may not always hold, the framework is essential to enable the estimation of conduction velocities in vivo and facilitates comparison with prior studies.

### White matter stimulation provides unique insight into velocity distributions

Previous studies estimating conduction velocity with intracranial measurements mostly focussed on estimating average velocities, rather than capturing full velocity distributions. Cortical surface recordings with electrocorticography (ECoG) across the arcuate fasciculus report typical latencies for the earliest negative N1 response in the range of ∼20-40 ms [52]. Using tractography to estimate the path length, these average latencies can be translated to conduction velocities of approximately 4 m/s [26,27]. While our estimates show some comparable average velocities for the arcuate fasciculus, we also observed many faster responses. These faster responses may be present in raw data reported in previous studies [26]. However, the larger variety of velocities may also be due to the fact that ECoG stimulates the cortical surface [53,57], whereas our white matter stimulation directly activates axons and triggers rapid antidromic spread. In contrast, cortical stimulation likely triggers more orthodromic responses with slower latencies due to synaptic delays at the measurement site. Overall, white matter stimulation may thus be better suited to capture velocity distributions.

Interestingly, Deep Brain Stimulation (DBS) studies often report evoked potentials with very short (<2 ms) to short (2–10 ms) latencies when stimulating the subthalamic nucleus (STN) and recording in motor, sensorimotor, and prefrontal cortical regions. These rapid cortical responses are thought to reflect propagation along the hyperdirect pathway, via antidromic transmission [58–60]. Assuming a distance of 10 cm from STN to cortex, these short latencies roughly correspond to speeds within a range of 10 to 50 m/s. This matches a small percent of our more rapid cortico-spinal tract estimates, while the majority of the distribution indicated slower velocities. It is possible that subthalamic stimulation mostly drives the thickest axons while missing smaller ones, either because they are too far away or because the stimulation current is below their activation threshold. This interpretation is consistent with well established evidence that thinner fibers have higher activation thresholds [61–63]. Moreover, computational estimates indicate that high-speed thalamocortical connections account for only ∼4% of the total white matter volume [64]. The short latencies found in subthalamic DBS may thus only capture a small portion of hyperdirect fibers, whereas the white matter stimulation used in our analyses likely drives a broader spectrum of fiber diameters.

### A long-tailed distribution in human white matter may support timing of brain function

We observed consistent differences in conduction velocities between bundles, however, variation was larger within bundles than between them. Beyond anatomical and functional factors, these results may be explained by the hierarchical temporal organization of neuronal activity [65], in which brain rhythms form a well structured system for spike activity within and between circuits over multiple temporal scales. This principle is hypothesized to be conserved across humans and other mammals, suggesting that the fundamental mechanisms of neural dynamics and temporal processing remain consistent despite the significant differences in brain volume. For this hypothesis to hold, structural compensations are required to maintain timing and rhythmic organization across species, with axon diameter and myelination playing particularly critical roles. These adaptations ensure that homologous connections between species reach their targets within approximately the same temporal windows. In this sense, if all axons in the brain were to uniformly compensate for brain size, the human brain would need to be substantially larger to accommodate enlarged axons. Instead, our results align with the presence of a long-tailed distribution of axon diameters, in which a small number of very fast, large-caliber axons are required to preserve global timing across the human brain.

### Differences between white matter bundles may support their function

Our results show reliable differences between white matter bundles. We hypothesize that these may be related to the distinct functional roles of each pathway. Association tracts such as the uncinate fasciculus, arcuate fasciculus, and the cingulum, exhibit conduction velocities that may be well suited to complex, integrative cognitive processing. The uncinate supports communication between the orbitofrontal cortex and the temporal lobe for socio-emotional and episodic memory [66], while the cingulum supports executive function and attention by linking cortical and subcortical sites in the limbic system [48]. Similarly, the arcuate fasciculus connects temporal and frontal regions to support cognitive functions such as high-level linguistic structures [67]. The conduction velocities within these bundles have to support multisensory integration and feedback processing. Because these functions require the integration of diverse information types, their timing must allow for more complex computation and synchronization across multiple cortical areas, rather than maximizing conduction speed.

In contrast, the significantly higher conduction velocities of the cortico-spinal tract are consistent with its role in the direct control of voluntary movement. The cortico-spinal tract originates in the motor and somatosensory cortices and descends through the internal capsule and brainstem to terminate on spinal motoneurons. Its high conduction velocity likely reflects the need for rapid signal propagation over long distances to enable precise motor execution [68]. This suggests that white matter structure may develop to support specific functions across the brain.

### Understanding conduction velocity to improve models of brain function

Several models of brain function use conduction velocity as a parameter. For example, assumed velocity ranges from 4 m/s in the virtual brain stimulation model [4], 5-10 m/s in models of resting-state fMRI connectivity [6,7], and to 5-26 m/s in neural mass models [8–10]. In this sense, instead of having single values, a parametric distribution function would be beneficial to encompass all these possible values and better describe brain function.

To exemplify how parametric distributions can be useful for more realistic modeling, we implemented them in a simple case: tractography visualization (Figure 7). Typically, streamlines are set to a uniform thickness. However, for a more realistic visualization, we can use the parametric distributions obtained for the conduction velocity of each bundle and sample values from the fitted parametric conduction velocity distributions for each bundle and use them to vary the thickness. Figure 7 shows how streamlines representing axons show high variability in thickness, which is more representative of biological reality.

**Figure 7.**
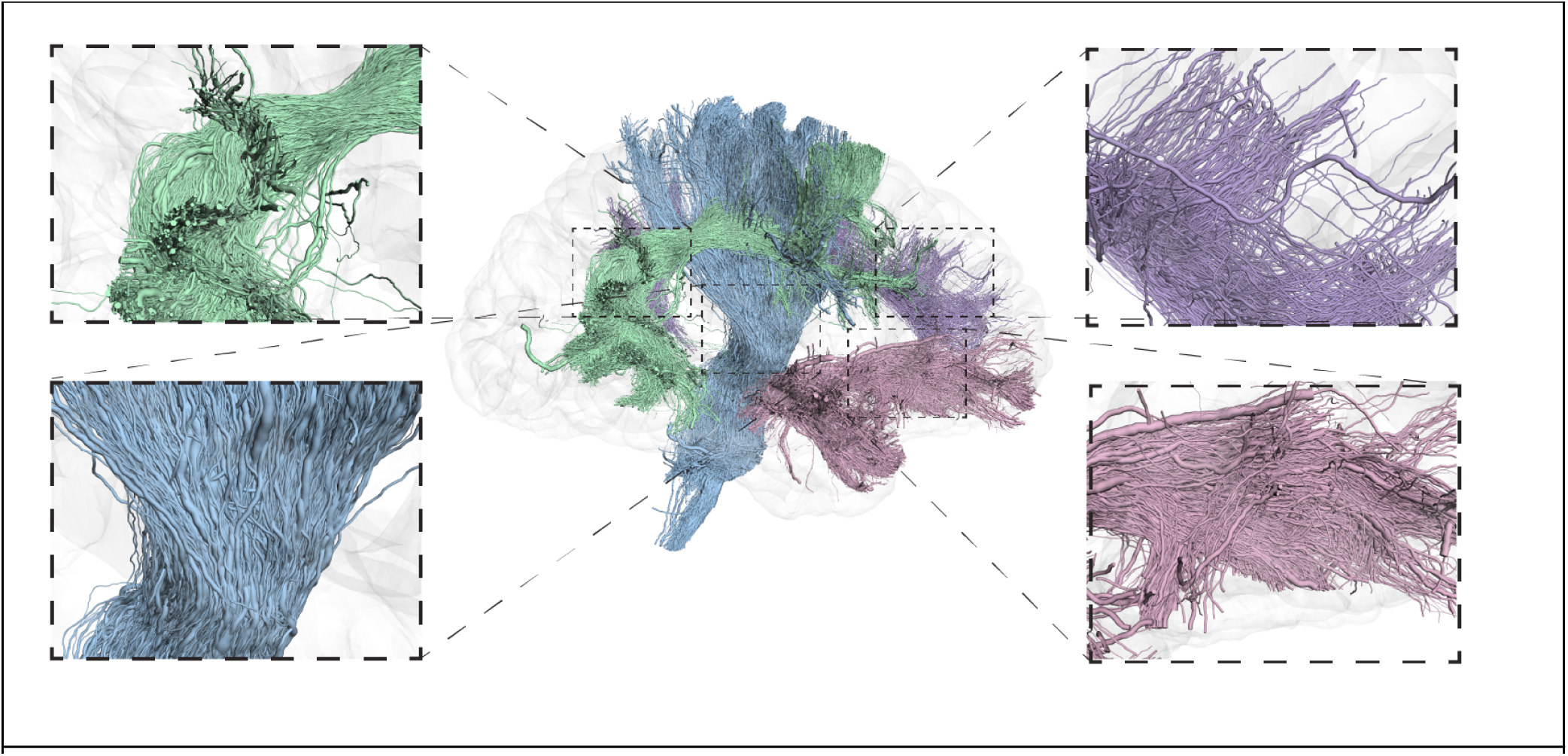
Streamline tractography visualization with varying diameters. Colors indicate bundle identity: arcuate fasciculus in green, cingulum in purple, uncinate fasciculus in pink, and cortico-spinal tract in blue. Streamline thickness, rather than color, represents the sampled axon diameter values. Streamline diameters were sampled from a bundle-specific, fitted log-normal distribution. The log-normal distribution was selected for the visualization, but we note that the GEV’s performed better in the cortico-spinal tract and the cingulum. This decision was based on the overall good fit of the log normal distribution, its parsimony, specifically its fewer parameters and simpler support structure compared to the GEV distribution. These sampled values were then translated into axon diameters through a linear relationship. Instead of setting a single thickness value for all streamlines, we used the sampled values to create more realistic visualizations of within-bundle variability in axon caliber.

## Conclusion

In summary, we introduce a novel pipeline that allows estimating distributions of axonal conduction velocity in the living human brain. By integrating sEEG stimulation with diffusion MRI tractography, this approach provides an unique in vivo framework for estimating conduction velocity along anatomically defined pathways. The resulting conduction velocity distributions show qualitative agreement with prior post-mortem histological observations of axon diameter distributions, showing skewness that reflects the presence of a majority of slower-conducting fibers and few exceptionally thick axons driving high-speed conduction. These distributions are key for understanding how structure supports the timing and integration of information in the human brain. By providing empirically derived conduction velocity distributions rather than single summary values, this approach adds a critical dimension for constraining and refining biophysically realistic models of large-scale human brain function.

## Methods

### Participants and Electrode Localization

Data were collected from neurosurgical patients with sEEG probes implanted during invasive epilepsy monitoring. A total of 17 patients provided informed consent to participate in the studies (ages 12-55, 9 female). The study was approved by the Mayo Clinic IRB (#15-006530 for iEEG and #19.007320 for MRI measurements).

Multicontact flexible sEEG probes (DIXI Medical) with electrode contacts 2 mm in length and 0.8 mm in diameter were implanted. The placement of the sEEG probes was selected by the clinical team for the purpose of seizure onset zone localization, with electrode coverage of different brain regions according to the individualizing clinical planning. Electrodes were localized using a computed tomography scan [69] and were coregistered to the T1-weighted (T1w) anatomic magnetic resonance imaging (MRI) using existing software ([70]; https://www.fil.ion.ucl.ac.uk/spm). MRI scans were autosegmented using FreeSurfer 7 ([71]; https://surfer.nmr.mgh.harvard.edu/). The segmentation was reviewed for accuracy, and sEEG electrodes were labeled according to the FreeSurfer Destrieux Atlas [72].

### MRI Scans

Participants were scanned on a Compact 3T head-size MRI system (GE Global Research, Niskayuna, New York, USA) equipped with a high-performance gradient system [73]. The high-performance gradient enables shorter EPI (echo planar imaging) echo spacing, which can reduce susceptibility-related geometric distortion commonly observed in conventional EPI acquisitions [74,75] This is particularly relevant for diffusion MRI, because tractography-based pathway length estimation depends on accurate spatial correspondence between anatomical and EPI images [74], and intracranial electrode locations.

High-resolution T1-weighted structural images were acquired using a 32-channel head coil. Sequence parameters were as follows: isotropic voxel of 0.8 mm, repetition time (TR) = 2491 ms, echo time (TE) = 3.1 ms, inversion time (TI) = 1000 ms, flip angle = 8° and an in-plane parallel imaging acceleration factor of 2.

T2-weighted images were acquired using a 32-channel head coil with a 3D fast spin-echo sequence (CUBE). Acquisition parameters included: isotropic voxel of 0.8 mm, TR = 3000 ms, TE = 72.5 ms, echo train length = 130, and flip angle = 90°. The sequence utilized fat saturation alongside parallel imaging with an acceleration factor of 2 in both the in-plane and through-plane directions.

To further minimize geometric distortion in the diffusion imaging, we used DIADEM (Distortion-free Imaging: A Double Encoding Method)-based diffusion imaging [76], a multi-shot EPI-based approach designed to reduce distortion and blurring relative to conventional single-shot EPI. Therefore, the conduction velocity estimates in this study were based on diffusion tractography derived from relatively distortion-minimized imaging, which is important for reliable estimation of white matter pathway distances. Subjects 01, 03-06, 09, 10, 12-15 and 17 were scanned with two series, each with three volumes at b= 0 s/mm2 and 48 directions at b = 1000 s/mm2, TR = 2659 ms; TE = 42.7 ms; TE_NE_ = 49.6 ms (TE_NE_ is navigator echo time for the DIADEM sequence) 78 slices at 1.8 mm thickness (zero slice gap), FOV of 216 x 216 mm, and acquisition matrix of 120 x 120 (interpolated to 240 x 240), yielding a final voxel size of 0.9 x 0.9 x 1.8 mm. Subjects 02, 07, 08, 11 and 16, were scanned with two series with each two volumes at b = 0 s/mm2 and 48 directions at b = 1000 s/mm2, TR = 2659 ms; TE = 42.7 ms; TE_NE_ = 49.6 ms, 70 slices were acquired at 2 mm thickness (zero slice gap), FOV of 216 x 216 mm, and acquisition matrix of 108 x 108, for a final voxel size of 2 x 2 x 2 mm.

### dMRI preprocessing

dMRI data were preprocessed to correct for subject motion and eddy currents and to align the dMRI images and T1w anatomic image using the Advanced Normalization Tools algorithm in QSIprep version 0.14.2 (Cieslak et al., 2021). Final images had a voxel size of 1.25mm^3^ in a matrix of 156 x 186 x 158. QSIprep includes denoising using the command “dwidenoise” in MRtrix3 (http://www.mrtrix.org) to improve the signal-to-noise ratio and to increase the accuracy of the diffusion parameter estimation and a B1 Bias Field Correction using the command “dwibiascorrect” in ANTs/MRtrix3 to correct for any inhomogeneity in the radio frequency field.

### Tractography and bundle segmentation

Whole-brain tractography (Figure 1-A) was performed using pyAFQ API [77]; https://tractometry.org/pyAFQ/reference/api/index.html). The Orientation Distribution Function (ODF) was solved using Constrained Spherical Deconvolution (CSD). Then, probabilistic tractography was performed using a local tracking approach. A whole brain mask was used as the seeding mask with a density of 2 seeds per voxel i. e. an array of 2×2×2 seeds for each voxel, being two for each direction. The maximum angle was set to 30 degrees, a step of 0.5mm was used with a minimum length of streamline of 20mm and a maximum of 250mm.

From the whole-brain tractography, AF, UF, Cing and CST were segmented using RecoBundles Algorithm [78]: a streamline-based registration and clustering model implemented in DIPY Python library [79].

### MRI metrics

The MRtrix software package (http://www.mrtrix.org) was used to estimate the diffusion tensor model from dMRI. Fractional anisotropy (FA) and radial diffusivity (RD) maps were subsequently derived.

The T1w/T2w ratio was computed using the Matlab MRTool package [45]. This pipeline standardizes the intensity histograms of the resulting ratio images, enabling robust across-subject statistical comparisons.

An electrode was considered to be in a bundle if it was located within a 4 mm radius of any streamline in the bundle. For a bundle to be considered in the study, at least one electrode from the stimulated pair and at least one recorded electrode must be located in the bundle. Additionally, at least one streamline must connect these two electrodes. The lengths of the streamlines (or segments of streamlines) between stimulated and recorded electrodes were measured from the closest point of the streamline to the stimulated electrode and the closest point of the streamline to the recorded electrode. We considered the conduction distance between the stimulated electrode pair and the recorded electrode as the shortest length among all streamlines (or segments of streamlines) connecting them (Figure 1-C). Finally, to quantify the FA, RD and T1w/T2w ratio associated with each electrode pair, the mean FA, the mean RD and the mean T1w/T2w ratio across all voxels intersected by the shortest streamline was computed.

### sEEG recordings and single pulse electrical stimulation

sEEG signals were measured on a g.HIamp biosignal amplifier (gTec) at 4800 Hz with up to 256 channels. After electrodes were implanted, seizures were monitored clinically. Once patients were back on medication, pairs of electrodes were stimulated with a single electrical pulse. Typical parameters were 6 mA, biphasic pulses of 100 μs width per phase. Stimulation amplitude was adjusted to 4mA in case the cortex was highly excitable (subjects 02, 11 and 13). Each pair was stimulated 10-12 times with ∼3 sec inter pulse interval with jitter (Figure 1-B). Seizure onset areas may affect later stimulation evoked responses, rather than the early evoked potentials of interest [80]. However, data were still carefully reviewed and excessive interictal activity or artifacts were marked and rejected in subsequent analyses. Moreover, Supplementary Table 4 shows that patients had highly variable seizure onsets, making it unlikely that the main results are driven by seizure networks. By design, dMRI also excludes tracking near electrodes on areas with anatomical abnormalities.

### sEEG Preprocessing

Recorded sEEG signals were preprocessed under a divergent paradigm, with stimulation delivered in one region and responses recorded across all other regions. Raw data were converted to a trial structure centered around the onset of electrical stimulation and corrected for DC offset. We considered one trial to be the response recorded after an individual SPES (from 0 to 1 s after the stimulation onset). We then ensured the data were free from reference contamination and re-referenced the data using an adjusted common average referencing (CAR) scheme [81,82]. We used this method only including channels with the lowest 10% variance as the common reference, which was shown to suppress noise while minimizing bias. Then, data was baseline corrected by subtracting the mean pre-stimulation amplitude for each trial from-100 to-10ms.

### Significance of responses

Significance and reliability of the responses were measured using the Canonical Response Parametrization method [83]. This algorithm allows measurement of the similarity and reliability across trials. In brief, for each electrode stimulated K times (K=10-x trials) and recorded through a T number of time points (over the interval 5ms ≤ t ≤ 1000ms for our particular case), a matrix *V*_(*TxK*)_ is built. V then is projected into each other trials 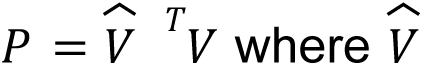 the unit-normalized V matrix. Matrix P is then sorted in a subset S (omitting self projections) with a total of *K*^2^ − *K* elements. The average over the set of cross-projections, *S̄*, summarizes the reliability of the response to stimulation. To assess how long there is a significant effect after stimulation, S can be constructed over different time periods to determine the duration of a reliable and statistically-meaningful response. This was accomplished by determining projection weights S and *S̄* as a function of time and quantifying a temporal profile, *S̄t*. As *S̄* may be thought as a reflection of mutual information between responses, the peak of *S̄t* represent the time past which further information is not reliably contained in the response. This peak time τ*_R_* is defined as the response duration. Significance of *S_TR_* against zero is assessed using a right tailed t-test (only half of the projections are considered to avoid artificial significance). The CRP method also gives an estimation of the canonical response C(t) (the characteristic shape of the BSEP), using a linear kernel-PCA (Principal Component Analysis) on the matrix V, being this the first component of the decomposition. Finally, individual trials can be parametrized in terms of C(t) as *V_k_*(*t*) = α*_k_C*(*t*) + ε*_k_*(*t*), where α is a constant scaling every trial and ε*_k_*(*t*) is the residual (containing noise and uncorrelated brain activity), with this description several metrics can be obtained for each trial, being of particular interest the “signal-to-noise” ratio SNR defined as 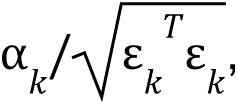 and a Coefficient of Determination (CoD) or *R*^2^, defined as 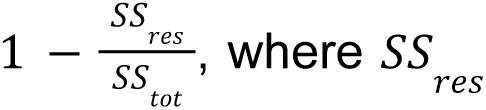 is the residual sum of squares and *SS_tot_* is the total sum of squares. We considered a response to be significant if it has a median CoD ≥ 0.10 and median SNR≥1 across trials. Also, we only took into account those signals with a p-value<0.05 when testing τ*_R_*.

### Latency measurement

Unlike ECoG, which relies on consistent N1 peaks for latency measurements, sEEG recordings are complicated by electrode laminar position, as this can cause polarity to change. Therefore, we defined conduction time as the latency of the first detectable peak following the stimulation artifact within a 1.8–40 ms window. Peak detection was performed using the *find_peaks* function from SciPy library 1.15.2 ([84]; https://docs.scipy.org/doc/scipy/reference/generated/scipy.signal.find_peaks.html). This function considers a peak as the sample whose two direct surrounding neighbors have smaller amplitude, however due to the noisy nature of sEEG signals this approach is not enough to accomplish the task. To improve robustness, we used the peak prominence as a filter, which is defined as the measurement of how much a peak stands out from the surrounding baseline of the signal and is defined as the vertical distance between the peak and its base (in μV for our signals, [85]). To ensure robust peak detection, an adaptive prominence was derived from the signal. This threshold was calculated from the pre-stimulus baseline period, spanning 100 to 10 ms prior to stimulus onset. To minimize the influence of signal outliers, the filter threshold was based on the Median Absolute Deviation (MAD, [86]) of the baseline samples, defined as follows:

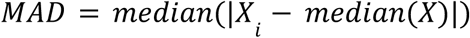

where X represents the set of baseline samples and X_i_ is the ith sample within X.

To mitigate the omission of early peaks in case of very noisy signals and ensure the capture of the earliest response, a ceiling value was implemented to prevent the threshold from becoming excessively high due to baseline fluctuations. Consequently, the final algorithm considers the earliest response of the signal as the first peak with a prominence of at least min(2*MAD,10μV). As the polarity of the signal on sEEG recordings is determined by its laminar position, responses were flipped to also find the first negative peak. Finally, we defined the conduction time as the shortest latency of the first positive or negative peak (Figure 1-D). To ensure data quality, we visually inspected the dataset and manually corrected identified errors. The overall distribution shapes remained consistent following these corrections and the removal of false positives. Furthermore, subsequent statistical analysis yielded unchanged results, maintaining statistically significant differences across all pairwise comparisons (Games-Howell test; Supplementary Fig. 1).

### Velocity computation

Conduction Velocity of the signal propagation between the stimulated electrode pair and the recorded electrode was computed as the conduction distance divided by the conduction time and reported in millimeters over milliseconds 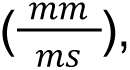 an example is shown in Figure 1-E.

### Axon Diameter Estimation

Axon diameter distributions were derived based on its reported linear relationship with conduction velocity [14]. In myelinated axons, this relationship with inner axon diameter has the following form:

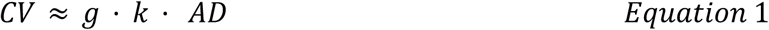

Where the value of inner axon diameter (AD) was derived from each conduction velocity (CV). The linear factor *k* has been approximated by several authors. In a seminal study Hursh [13] established a constant value of approximately k = 6 for feline peripheral nerves. Subsequent investigations incorporating detailed morphological features, such as internodal length, proposed a value of 5.5 [15,87]. Conversely, other experimental data have suggested slightly different values starting from 4 [88] to around 8 [89,90]. In order to account for this variability and encompass the broad range of reported literature values, we used scaling factors of 4, 6, and 8 for our analysis. While previous studies in rodents(Knowles et al. 2022; Chau Loo Kung et al. 2023) and humans (Kim et al. 2026) have shown small but statistically significant lower mean g-ratios in epileptic networks (indicating a thicker myelin sheath relative to the inner axon diameter) due to aberrant oligodendrocyte precursor cell proliferation, these differences are very small (<0.06) in all cases. Therefore, for our estimations, a constant g-ratio (defined as the ratio between the inner and outer fiber diameters) of 0.7 was assumed [37,38], as illustrated in Figure 5.

## Statistical Analysis

### Hemisphere Comparison

To assess interhemispheric differences in white matter bundles, we compared group medians using two-sided non-parametric permutation tests with 10,000 resamples [33]. This approach was selected to provide exact p-values because it doesn’t assume normality in the underlying velocity distributions. To maintain a family-wise error rate of α=0.05 across the four homologous bundle pairs, p-values were adjusted using the Bonferroni correction (Dunn, 1961). Statistical analyses were implemented in Python using the SciPy library (version 1.15.2; Virtanen et al., 2020).

### Pairwise Comparison

To assess differences in mean velocity across multiple bundles, we employed Welch’s ANOVA [91], which is robust to violations of the assumption of equal variances. In the event of a significant main effect, post-hoc pairwise comparisons were conducted using the Games-Howell test [92] to account for both heterogeneity of variance and unequal sample sizes. Statistical analyses were performed using the Pingouin library 0.5.5 ([93]; https://pingouin-stats.org/build/html/index.html)

### Parametric distribution fitting

For each bundle, Log-normal, Gamma and Generalized Extreme Value density distribution functions were fitted.

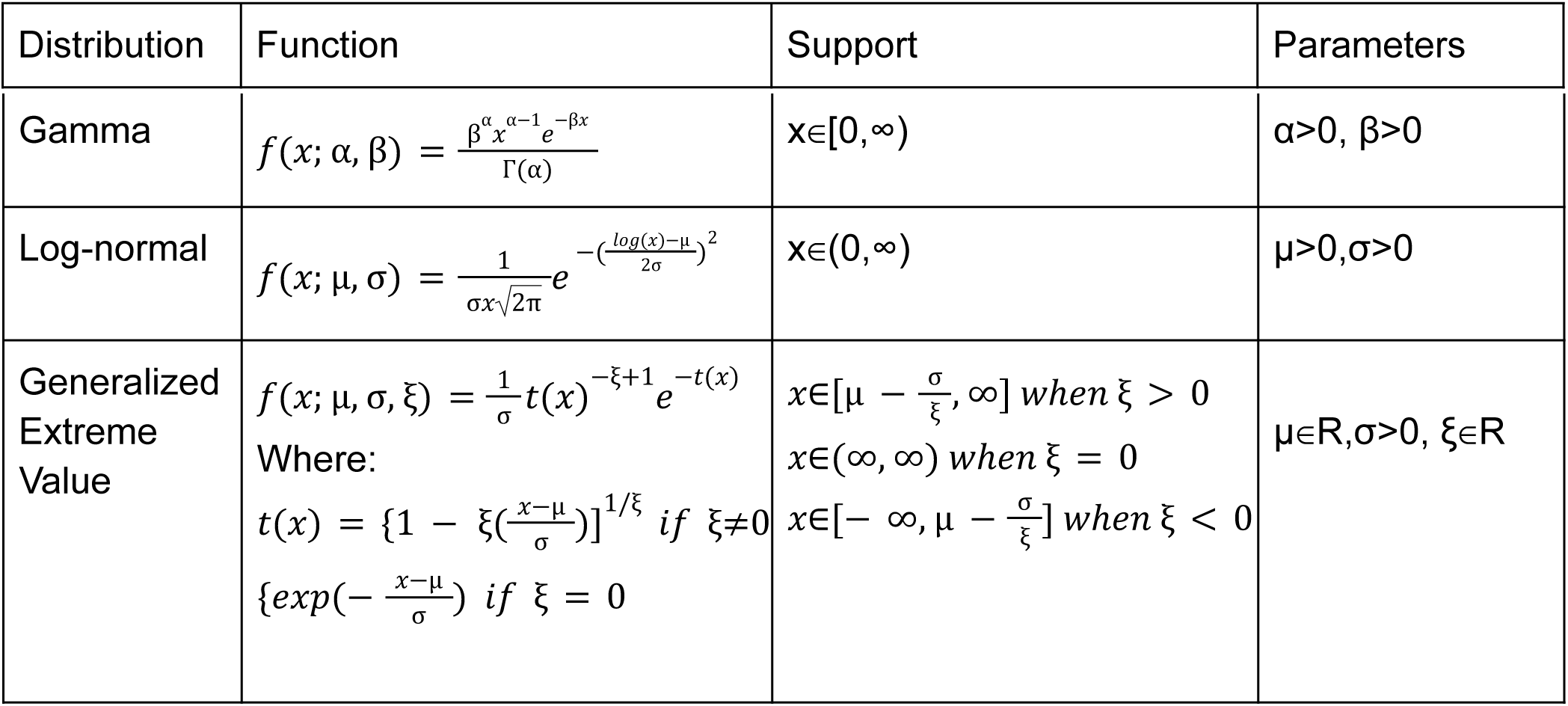

The Log-likelihood (*l*), Akaike Information Criterion (AIC) and Bayesian Information Criterion (BIC) were computed as follows:

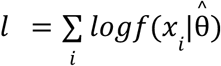

Where *f*(*x_i_*|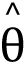) is the corresponding distribution function for point *x_i_* using the estimated 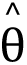 parameters.

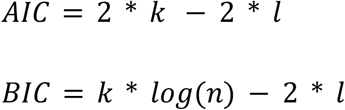

Where k is the number of the estimated parameters, (k=2 for Log-normal and Gamma distributions and k=3 for Generalized Extreme Value distribution) and n is the number of observations in a given bundle. The SciPy library 1.15.2 ([84]; https://docs.scipy.org/doc/scipy/index.html) was used to fit the three distributions, calculate the Log-likehood, the AIC and the BIC.

### Linear Mixed-Effects Model

Since the data involved multiple subjects and varying sample sizes per bundle, a standard linear regression would violate the assumption of independence (samples from the same subject are more alike than samples from different subjects). Hence, we used a Linear Mixed-Effects Model with the MRI metric as the fixed effect and the subject as the random effect as follows:

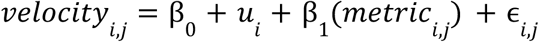

Where *i* represents the ith individual subject (the grouping factor), *j* is the jth observation within patient, *velocity_i_*_,*j*_ is the dependent variable, β_0_ is the global intercept, *u_i_* is the random intercept for each subject, assuming that some people may have higher velocity baselines, *metric_i_*_,*j*_ is the primary predictor representing the MRI metric (FA, RD or T1w/T2w), β_1_ is the fixed effect slope for the MRI metric and ɛ*_i_*_,*j*_ the residual error of the model. Statsmodels library 0.14.5 ([94]; https://www.statsmodels.org/stable/index.html) was used to perform the Linear Mixed Effects regression.

## Data availability

Code for sEEG preprocessing and response significance was written in Matlab R2024a. Code for tractography, bundle segmentation, latency, figures and analysis was written in Python 3.12.9. Code and data to reproduce the main results will be made available on github and OpenNeuro upon publication.

## Acknowledgements

The authors would like to thank the patients who participated in this study, Karla Crocket, Cindy Nelson and other staff and colleagues who assisted at Saint Mary’s Hospital. This work was supported by the National Institutes of Health (NIH) grants R01-EY35533 (to D.H) and K23-NS136792 (to N.G). The content is solely the responsibility of the authors and does not represent the official views of the NIH.

