## Supplemental Material for "White matter conduction in the human brain is mostly slow, with rare high velocity connections"

**Supplementary Material**

| **Bundle** | **Number of significant responses (n)** | **Mean (m/s)** | **SD**  **(m/s)** | **Median**  **(m/s)** | **Skewness** | **Max. (m/s)** | **Min.**  **(m/s)** |
| --- | --- | --- | --- | --- | --- | --- | --- |
| Right Uncinate Fasciculus | 138 | 5.683 | 4.168 | 4.168 | 2.301 | 30.080 | 0.774 |
| Left Uncinate Fasciculus | 131 | 7.251 | 5.884 | 5.142 | 0.774 | 24.449 | 0.853 |
| Right Cingulum | 431 | 7.975 | 4.415 | 6.833 | 1.672 | 35.142 | 1.007 |
| Left Cingulum | 162 | 8.330 | 4.047 | 7.251 | 1.158 | 23.466 | 1.062 |
| Right Arcuate Fasciculus | 625 | 10.239 | 7.483 | 8.123 | 1.586 | 40.199 | 0.966 |
| Left Arcuate Fasciculus | 265 | 8.742 | 5.516 | 7.399 | 1.158 | 33.600 | 0.992 |
| Right Cortico-Spinal Tract | 825 | 13.433 | 6.876 | 12.266 | 0.831 | 37.600 | 1.292 |
| Left Cortico-Spinal Tract | 82 | 9.905 | 4.500 | 9.357 | 0.313 | 21.600 | 1.183 |
| **Supplementary Table 1.** Statistical descriptors of computed velocities for all different bundles and hemispheres. | | | | | | | |

| **Bundle** | **Fitted Distribution** | **Estimated Parameters** | **Log-likelihood** | **AIC** | **BIC** |
| --- | --- | --- | --- | --- | --- |
| Arcuate Fasciculus | Log-Normal | μ=2.0503,σ=0.6956 | **-2764.6954** | **5533.3909** | **5542.9734** |
|  | Gamma | α=2.3134, β=0.2362 | -2777.7806 | 5559.5613 | 5569.14382 |
|  | GEV | μ=6.2998,σ=4.0166, ξ=-0.2484 | -2769.2124 | 5544.4248 | 5558.7985 |
| Uncinate Fasciculus | Log-Normal | μ=1.5242,σ=0.8466 | **-746.9524** | **1497.9049** | **1505.0944** |
|  | Gamma | α=1.6201, β=0.2513 | -753.8812 | 1511.7625 | 1518.9520 |
|  | GEV | μ=3.3723,σ=2.5939, ξ=-0.4909 | -754.0385 | 1514.0771 | 1524.8612 |
| Cingulum | Log-Normal | μ=1.9527,σ=0.5377 | -1631.5146 | 3267.0292 | 3275.7995 |
|  | Gamma | α=3.8444, β=0.4762 | -1625.7407 | 3255.4815 | 3264.2519 |
|  | GEV | μ=6.0997,σ=3.0538, ξ=-0.0648 | **-1619.4253** | **3244.8507** | **3258.0063** |
| Cortico-Spinal Tract | Log-Normal | μ=2.4237,σ=0.5918 | -3009.5309 | 6023.0619 | 6032.6822 |
|  | Gamma | α=3.4911, β=0.2661 | -2961.4111 | 5926.8223 | 5936.4426 |
|  | GEV | μ=10.0778,σ=5.4400, ξ=0.0225 | **-2957.5281** | **5921.0562** | **5935.4866** |
| **Supplementary Table 2. Results of fitted parametric distributions.** Estimated parameters for the log-normal, gamma, and generalized extreme value (GEV) distributions are reported, together with model fit indices, including log-likelihood, Akaike Information Criterion (AIC), and Bayesian Information Criterion (BIC). | | | | | |

| Bundle | MRI Metric | Parameter | Coefficient | SE | Z-statistic | p-value | Confidence Intervals [0.025,0.975] |
| --- | --- | --- | --- | --- | --- | --- | --- |
| UF | FA | Intercept | -1.764 | 2.671 | -0.661 | 0.509 | [-6.998, 3.470] |
|  |  | Slope | 19.199 | 6.255 | 3.069 | **0.002** | [6.938, 31.459] |
|  | RD | Intercept | 19.247 | 4.175 | 4.610 | 0.000 | [11.063, 27.431] |
|  |  | Slope | -19958.800 | 6331.666 | -3.152 | **0.002** | [-32368.637, -7548.963] |
|  | T1w/T2w | Intercept | 5.623 | 2.966 | 1.896 | 0.058 | [-0.191, 11.437] |
|  |  | Slope | 0.358 | 3.704 | 0.097 | 0.923 | [-6.901, 7.617] |
| Cing | FA | Intercept | 1.630 | 0.151 | 10.810 | ~0.0 | [1.334, 1.925] |
|  |  | Slope | 0.705 | 0.323 | 2.182 | **0.029** | [0.072, 1.337] |
|  | RD | Intercept | 2.340 | 0.229 | 10.213 | ~0.0 | [1.891, 2.789] |
|  |  | Slope | -651.989 | 362.688 | -1.798 | 0.072 | [-1362.844, 58.865] |
|  | T1w/T2w | Intercept | 3.719 | 1.916 | 1.941 | 0.052 | [-0.036, 7.474] |
|  |  | Slope | 5.321 | 2.363 | 2.252 | **0.024** | [0.691, 9.952] |
| AF | FA | Intercept | 5.150 | 1.959 | 2.628 | 0.009 | [1.310, 8.990] |
|  |  | Slope | 10.927 | 4.520 | 2.417 | **0.016** | [2.067, 19.787] |
|  | RD | Intercept | 24.623 | 3.129 | 7.868 | ~0.0 | [18.489, 30.756] |
|  |  | Slope | -24095.925 | 4890.276 | -4.927 | **~0.0** | [-33680.689, -14511.160] |
|  | T1w/T2w | Intercept | 4.741 | 2.074 | 2.286 | 0.022 | [0.677, 8.805] |
|  |  | Slope | 5.707 | 2.306 | 2.475 | **0.013** | [1.188, 10.226] |
| CST | FA | Intercept | 6.576 | 1.778 | 3.698 | ~0.0 | [3.090, 10.061] |
|  |  | Slope | 12.169 | 3.985 | 3.054 | **0.002** | [4.358 19.979] |
|  | RD | Intercept | 17.933 | 3.168 | 5.660 | ~0.0 | [11.723, 24.143] |
|  |  | Slope | -10263.261 | 5135.137 | -1.999 | **0.046** | [-20327.943, -198.578] |
|  | T1w/T2w | Intercept | 9.391 | 2.219 | 4.232 | 0.000 | [5.042, 13.740] |
|  |  | Slope | 2.867 | 2.624 | 1.093 | 0.275 | [-2.276, 8.011] |
| **Supplementary Table 3. Linear mixed-effects model results.** Estimated parameters from the fitted linear mixed-effects models are reported for each of the four white matter bundles and for both DTI metrics. Statistically significant slope effects (*p* < 0.05) are indicated in bold. | | | | | | | |

| **Subject** | **Sex** | **Age** | **Hemisphere Implanted** | **Bundle Coverage** | **Seizure Onset Zones (Destrieux labels)** |
| --- | --- | --- | --- | --- | --- |
| Sub-01 | Male | 31 | Bilateral | AFL,AFR,UFL,CSTL,CingL | Left_Hippocampus, lh_S_temporal_sup, lh_G_temp_sup-Lateral, lh_S_circular_insula_inf, lh_G_temp_sup-G_T_transv, lh_Pole_occipital, lh_S_oc_middle_and_Lunatus , lh_S_oc-temp_med_and_Lingual, ,lh_S_oc-temp_lat |
| Sub-02 | Male | 35 | Bilateral | AFL,UFL,CingL | Left_Hippocampus, lh_S_temporal_sup, lh_G_temp_sup-Lateral, lh_G_temporal_middle, Right_Amygdala, Right_Hippocampus, rh_S_temporal_sup, rh_G_temporal_middle |
| Sub-03 | Female | 16 | Bilateral | AFR,UFR,CSTR,CingR | rh_G_front_sup, rh_S_precentral-sup-part, rh_G_precentral, rh_S_circular_insula_inf, rh_G_temp_sup-Plan_polar, rh_G_temp_sup-G_T_transv, rh_G_temp_sup-Lateral, rh_S_temporal_sup, rh_G_temp_sup-Lateral |
| Sub-04 | Female | 12 | Bilateral | AFL,UFL | Left_Hipocampus, lh_S_temporal_sup, lh_G_temp_sup-Plan_tempo, lh_G_pariet_inf-Supramar, lh_S_circular_insula_inf, lh_G_Ins_lg_and_S_cent_ins, lh_G_insular_short, lh_S_circular_insula_sup |
| Sub-05 | Female | 54 | Bilateral | UFR | lh_S_temporal_sup, lh_G_temporal_middle, lh_G_temp_sup-Plan_tempo, lh_G_temp_sup-Lateral, lh_Pole_temporal |
| Sub-06 | Female | 21 | Bilateral | AFL,CSTL,CingL | rh_G_front_sup, rh_S_front_middle, rh_S_temporal_sup, rh_G_pariet_inf-Supramar, lh_S_front_middle, lh_G_rectus, lh_G_front_middle, lh_G_front_sup, lh_S_front_middle, lh_S_front_sup, lh_S_front_inf,lh_G_postcentral, lh_G_pariet_inf-Supramar |
| Sub-07 | Female | 16 | Right | AFR,UFR,CSTR,CingR | rh_S_central, rh_G_postcentral, Right_Thalamus, rh_G_and_S_paracentral, rh_G_front_sup, rh_G_and_S_cingul-Mid-Post, rh_S_precentral-sup-part, rh_G_precentral |
| Sub-08 | Female | 55 | Bilateral | AFR,UFR,CSTR,CingR | Right_Hippocampus, rh_S_oc-temp_med_and_Lingual, rh_G_temporal_middle |
| Sub-09 | Female | 18 | Bilateral | AFL,CSTL | lh_G_occipital_middle, lh_S_oc-temp_lat, rh_S_oc_middle_and_Lunatus |
| Sub-10 | Male | 18 | Left | AFL,UFL,CSTL,CingL | lh_Pole_temporal, lh_G_temporal_middle |
| Sub-11 | Male | 35 | Right | AFR,CSTR | rh_G_temp_sup-Lateral, rh_G_insular_short, rh_S_circular_insula_sup, rh_G_Ins_lg_and_S_cent_ins, rh_G_insular_short, rh_S_circular_insula_inf |
| Sub-12 | Male | 16 | Right | AFR,UFR,CSTR,CingR | Right_Thalamus, Right_Hippocampus, rh_Lat_Fis-post, rh_G_temp_sup-Plan_tempo,rh_S_circular_insula_inf, rh_Lat_Fis-post, rh_G_and_S_subcentral, rh_G_pariet_inf-Supramar |
| Sub-13 | Male | 27 | Bilateral | AFR,CSTR | rh_Pole_temporal, rh_G_oc-temp_lat-fusifor, ,rh_S_oc-temp_med_and_Lingual, rh_S_collat_transv_ant, rh_S_oc-temp_lat, rh_G_temporal_inf |
| Sub-14 | Female | 15 | Right | AFR,CSTR | rh_S_postcentral, rh_S_central |
| Sub-15 | Male | 43 | Bilateral | AFR,UFR,CSTR,CingR | rh_G_front_middle |
| Sub-16 | Male | 19 | Bilateral | AFL | Right_Putamen |
| Sub-17 | Female | 43 | Left | AFL,UFL,CSTL,CingL | Left_Hippocampus, Left_Inf_Lat_Vent, Left_Amygdala |
| **Supplementary Table 4**. Summary of participants for this work, electrode coverage and seizure onset zones | | | | | |

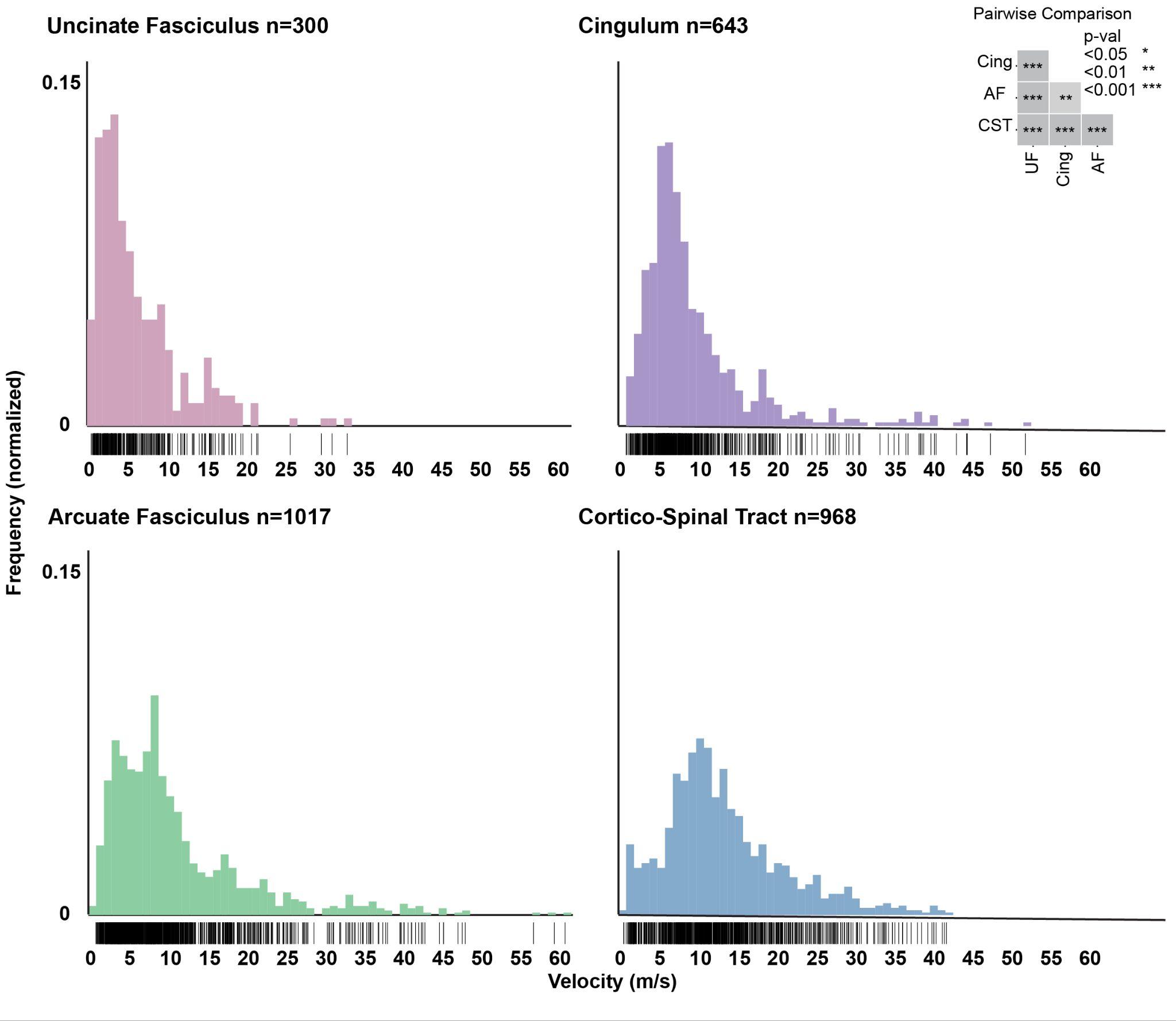

**Supplementary Figure 1. Histograms of pooled uncleaned data for four major white matter bundles.** Pooled uncleaned data of all subjects and hemispheres for Uncinate Fasciculus (pink), Arcuate Fasciculus (green), Cingulum (purple) and Cortico-Spinal Tract (blue). Individual observations are displayed below the histograms as black lines. Pairwise comparisons via Games-Howell test revealed significant differences between all bundles; AF:CST (p < 0.001), AF:Cing (p < 0.01), AF:UF (p < 0.001), CST:Cing (p < 0.001), CST:UF (p < 0.001), and UF:Cing (p < 0.001).
